# Multiple selection signatures in farmed Atlantic salmon adapted to different environments across Hemispheres

**DOI:** 10.1101/525550

**Authors:** M.E. López, T. Linderoth, A. Norris, J.P. Lhorente, R. Neira, J.M. Yáñez

## Abstract

Domestication of Atlantic salmon started approximately forty years ago, using both artificial and natural selection strategies. Such selection methods are likely to have imposed distinctive selection signatures on the salmon genome. Therefore, identifying differences in selection signatures may give insights into the mechanism of selection and candidate genes of biological and productive interest. Here, we used two complementary haplotype-based statistics, the within-population integrated Haplotype Score test (|iHS|) and the cross-population Extended Haplotype Homozygosity test (XP-EHH) to compare selection signatures in four populations of Atlantic salmon with a common genetic origin. Using |iHS| we found 24, 14, 16 and 26 genomic regions under selection in Pop-A, Pop-B, Pop-C, and Pop-D, respectively. While using the XP-EHH test we identified 27, 25 and 15 potential selection regions in Pop-A/Pop-B, Pop-A/Pop-C and Pop-A/Pop-D, respectively. These genomic regions harbor important genes such *igf1r* and *sh3rf1* which have been associated with growth related traits in other species. Our results contribute to the detection of candidate genes of interest and help to understand the evolutionary and biological mechanisms for controlling complex traits under selection in Atlantic salmon.

## 2. BACKGROUND

Atlantic salmon (*Salmo salar* L) were first farmed in Norway during the 1960s, and have now become one of the most important aquaculture species (FAO 2016). Despite a generation interval of three to four years, breeding programs have achieved rapid improvement of economically important traits such as growth, sexual maturation and disease resistance (Gjedrem *et al*. 2012). One of the first farmed populations named Mowi strain, was established with fish from west coast rivers in Norway, with major contributions from River Bolstad in the Vosso watercourse, River Årøy and possibly from the Maurangerfjord area (Verspoor *et al*. 2007). Salmon from the Vosso and Årøy rivers are characterized by large size and late maturity (Verspoor *et al*. 2007). Phenotypic selection for growth, late maturation and fillet quality was the focus in this population until 1999 (Glover *et al*. 2009). Ova from this population were imported into Fanad Peninsula, Ireland between 1982 and 1986 to establish an Irish-farmed population (Norris 1999). Similarly, ova from this Irish farmed population were introduced to Chile in the early 1990s. These stocks were subsequently adapted to the biotic and abiotic factors present in southern hemisphere conditions. Artificial selection and adaptation to captive environments has left detectable genomic patterns in farmed Atlantic salmon populations, as evidenced by differences between wild and farmed populations for several traits, such as growth rate (Thodesen *et al*. 1999; Glover *et al*. 2009; Solberg *et al*. 2012), predator awareness (Einum and Fleming 1997) and gene transcription patterns (Roberge *et al*. 2006; Bicskei *et al*. 2014; CHristie *et al*. 2016).

Domestication processes are likely to have exerted selection pressures on certain genomic regions that underlie traits of human interest or other traits involved in adaptation to captive environments. Accordingly, positive selection pressures will cause the frequency of alleles underlying favorable traits to increase rapidly in these domesticated populations. Linkage disequilibrium between favorable mutations and neighboring loci will increase and spread, given there is little opportunity for recombination over the brief time since the onset of intense selection (Sabeti *et al*. 2002). Analyses of these selection signatures in domestic animals can provide further insights into the genetic basis of adaptation to diverse environments and genotype/phenotype relationships (Oleksyk *et al*. 2010; Andersson 2012). Access to genomic data through next-generation sequencing and high-throughput genotyping technologies have made the comparison of genomic patterns of SNP variation between different livestock breeds possible, allowing for the identification of putative genomic regions and genes under selection in various species including cattle (Flori *et al*. 2009), horses (Petersen *et al*. 2013; Frischknecht *et al*. 2016), sheep (Kijas *et al*. 2012; Fariello *et al*. 2014), pigs (Amaral *et al*. 2011), Atlantic salmon (Vasemägi *et al*. 2005; Vasemägi *et al*. 2012; Mäkinen *et al*. 2014; Gutierrez *et al*. 2015; López *et al*. 2018) and tilapia (Hong Xia *et al*. 2015).

There are several approaches for detecting selection signatures in the genome, one of which relies on the length or variability of haplotypes. Directional selection acting on a new beneficial mutation results in the haplotype harboring the mutation to increase in frequency and to be longer than average. In order to exploit this, Sabeti et al (2002), proposed the extended haplotype homozygosity (EHH) statistic to detect of positive selection in a population, which is specifically the probability that two randomly selected haplotypes are identical-by-descent over their entire length around a core SNP (Sabeti et al 2002). This concept forms the basis for other haplotype homozygosity based metrics, such as the relative EHH (REHH) (Sabeti *et al*. 2002) and the widely-used integrated Haplotype Score (|iHS|) (Voight *et al*. 2006). |iHS| compares EHH between derived and ancestral alleles within a population and has the most power to detect selection when the selected allele is at intermediate frequencies in the population (Sabeti *et al*. 2006; Voight *et al*. 2006). To detect selection signatures between populations, the cross-population Extended Haplotype Homozygosity test (XP-EHH) compares the integrated EHH profiles between two populations at the same SNP. It was designed to detect ongoing or nearly fixed sites harboring selection in one population (Sabeti *et al*. 2007).

Although previous studies have already been carried out to detect selection signatures in Atlantic salmon (MÄKinen *et al*. 2014; Gutierrez *et al*. 2015; Liu *et al*. 2016; López *et al*. 2018) exploration of selection signatures in additional populations will illuminate how genetic variation among the different strains, adapted to different culture conditions, across hemispheres has not been assessed yet. Herein we used an Affymetrix 200K SNP array dataset to investigate selection signatures in farmed Atlantic salmon populations from the same origin, cultivated in Ireland and Chile. We identified several selection signatures using two haplotype-based approaches (|iHS| and XPEHH) at the whole genome level in four Atlantic salmon populations. These findings are important as they highlight regions of the genome that might benefit economically relevant attributes, such as growth, resistance to local diseases and adaptation to specific environmental conditions.

## 3. MATERIALS AND METHODS

### Samples, genotyping and quality control

We used a total of 270 individuals from four farmed Atlantic salmon populations of Norwegian origin (Pop-A, n = 40; Pop-B, n = 71; Pop-C, n = 85; Pop-D, n = 74). Pop-A fish are from the Irish strain (Fanad) originating from the west coast Rivers of Norway, as described in the Introduction section. Artificial selection for improving growth, maturity and fillet quality was applied from the beginning in this population (Glover *et al*. 2009). We estimated that this population had been under artificial selection for at least ten generations. Pop-B and Pop-C are two different Chilean populations, established with fish from two different year classes of the same Irish strain (Fanad) in the 1990s. Pop-B and Pop-C have been farmed and adapted to the Los Lagos Region, Chile (42°S 72°O). Pop-D is another Chilean population founded with fish from the same Irish farmed strain but adapted to the XII^nd^ Region, Magallanes, Chile (53°S 70°O). Pop-B, Pop-C and Pop-D populations experienced four generations of selective breeding for growth in Chilean farming conditions at the time of sampling.

Genotyping of all populations was performed using Affymetrix’s Atlantic salmon 200K SNP Chip described in YáñEZ *et al*. (2016). We assessed SNP quality control using Axiom Genotyping Console (GTC, Affymetrix) and SNPolisher (an R package developed by Affymetrix) i) removing SNPs that did not match with high quality clustering patterns, according to the best practices recommended by Affymetrix, *ii*) removing SNPs with call rate lower than 95% and iii) we discarded individuals with genotyping call rate under 90%. We used only SNPs that mapped to chromosomes in the newest version of the Atlantic salmon reference genome, ICSAG_v2 (GenBank: GCA_000233375.4). After quality control filtering, 146,102 SNPs remained for downstream analyses.

### Genetic diversity and population structure

We evaluated genetic diversity in terms of the observed heterozygosity (H_O_) and expected heterozygosity (H_E_) calculated with PLINK v1.07 (Purcell *et al*. 2007). To investigate population structure based on individual ancestry proportions, we performed model-based clustering assuming no prior knowledge about strain origins in ADMIXTURE 1.2.2 (Alexander *et al*. 2009). We performed 10 separate randomly seeded runs for each number K of ancestral populations (1<K<20) and selected the optimum K according to the lowest value of the cross-validation error. The aforementioned analyses were conducted using a total of 20,000 SNPs after retaining only those with linkage disequilibrium (LD) values of at most 0.2 to minimize possible confounding effects of LD on the underlying patterns of genetic structure.

### Selection signatures, gene annotation and functional analyses

To detect potentially regions harboring selection signatures, two complementary haplotype-based detection methods, iHS and *XPEHH*, were used for within and between population analyses, respectively.

### Detection of within-population selection signatures using *iHS*

The iHS score is based on the ratio of extended haplotype homozygosity (EHH) for haplotypes anchored with the ancestral versus derived allele. The ancestral allele state for salmon is unknown and so to avoid losing SNPs by trying to polarize them from publicly available outgroup references, we assumed that the major allele represented the ancestral state as used by Bahbahani *et al* (2015). We phased the haplotypes using Beagle (Browning and Browning 2009). Singlesite iHS values were calculated across the genome for each population. |iHS| scores were calculated using the REHH package (Gautier and Vitalis 2012) and a score threshold of 3.0 was used to infer candidate genomic regions under selection.

### Detection of between-population selection signatures using *XP-EHH*

The XP-EHH statistic compares the integrated EHH between two populations at the same SNP, in order to identify selection based on overrepresented haplotypes in one of the populations, detecting entirely or approximately fixed sites (Sabeti *et al*. 2007). The direction of selection can be determined from the sign of XP-EHH scores, whereby negative XP-EHH scores suggest selection in the ‘reference’ population, whereas positive scores suggest selection in the ‘observed’ population. Pop-A was used as the reference population to the other three populations, hence there were three pairs of comparisons.

### Gene functional annotation

Genomic regions harboring SNPs showing evidence of selection were annotated based on the ICSAG_v2 reference genome (Lien *et al*. 2016) using SnpEff (Cingolani *et al*. 2012). Gene transcripts from these candidate regions were aligned (using blastx) (Altschul *et al*. 1990) to the zebra fish *(Danio rerio)* peptide reference database (downloaded from http://www.ensembl.org/) to determine gene identify. As evidence of homology we used an e-value □ 0 and then retrieved the zebra fish gene identifiers and gene ontology (GO) information from the ensembl biomart database (http://www.ensembl.org/biomart).

## 4. RESULTS

### Genetic diversity and structure

We investigated genetic diversity within each population using SNPs filtered for missing data per individual (max 10%), missing data per marker (max 5%) and allele frequency (min 5%) as described in the Materials and Methods section. A total of 146,103 SNPs were retained for analyses after these quality control steps. Observed heterozygosity levels were similar across the four domestic populations. And was slightly higher than expected for populations A, B and C, and even higher in population D (See Table 1).

**Table 1.** Mean genetic diversity (Observed heterozygosity and expected heterozygosity) of four Atlantic salmon populations

| Population | H<sub>o</sub> | H<sub>e</sub> |
| --- | --- | --- |
| Pop-A | 0.38±0.16 | 0.37±0.15 |
| Pop-B | 0.40±0.15 | 0.39±0.14 |
| Pop-C | 0.40±0.14 | 0.39±0.13 |
| Pop-D | 0.46±0.22 | 0.37±0.16 |

Admixture analysis was used to determine the composition of ancestral lineages among individuals to offer insight into the observed genetic variation. We found K=12 ancestral lineages to be optimal in describing the ancestry of the individuals across the 4 populations (Figure 1).

**Figure 1.**
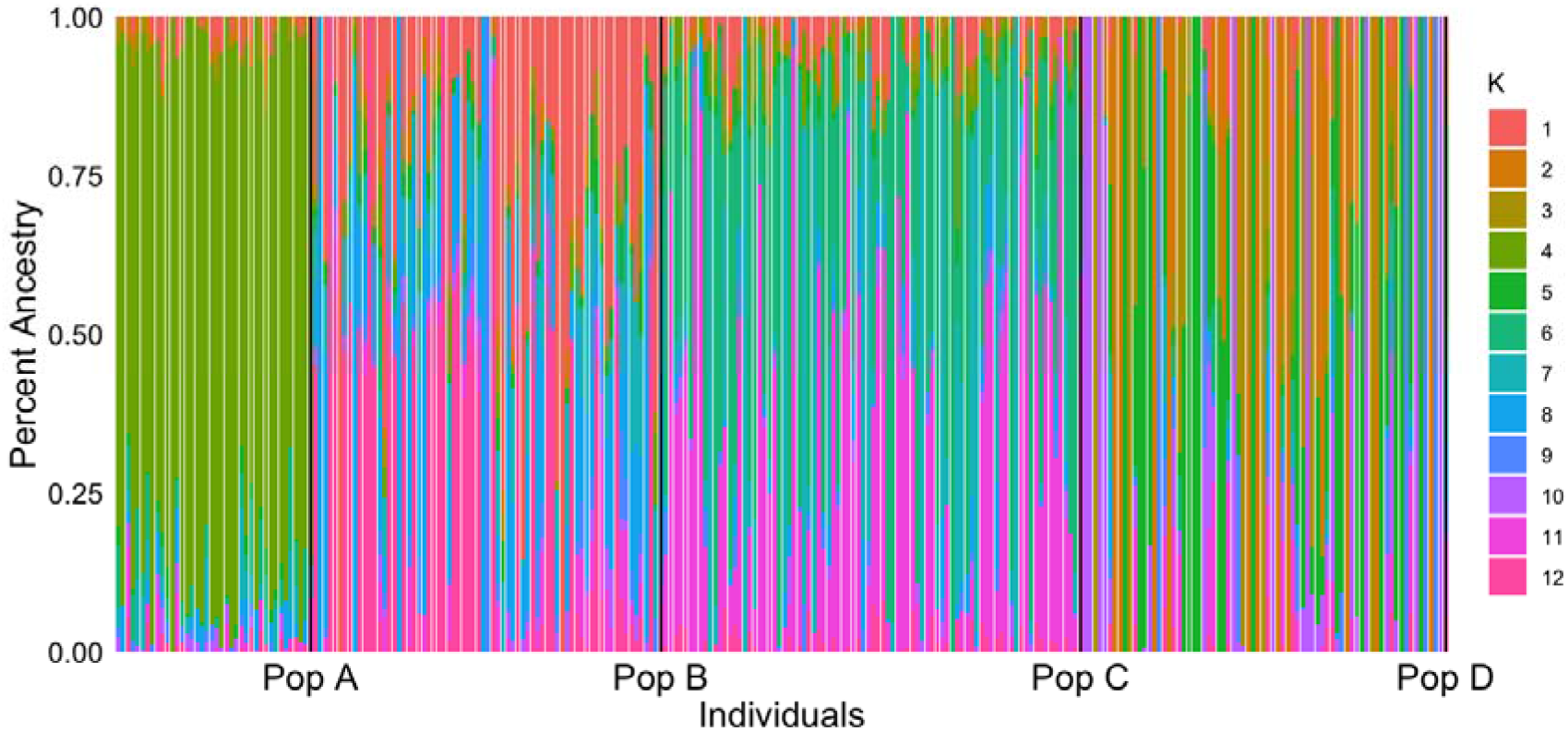
Individual assignment probabilities generated with ADMIXTURE (1·K·12). Each color represents a cluster, and the ratio of vertical lines is proportional to assignment probability of and individual to each cluster.

### Candidate regions under selection - |iHS|

We used the haplotype-based |iHS| test to look for selection within populations. For each population we defined candidate selection regions using the thresholds of |iHS| > 3 (Figure 2 and Table 2). Candidate regions were retained if two SNPs separated by ≤ 500 Kb passed this threshold and were annotated using the positions of the first and last SNP as boundaries, extending 500 Kb to each side. In Pop-A we identified 120 markers putatively under selection among ten chromosomes, Ssa02 and Ssa10 combined had approximately 60 SNPs. The highest score (-log(p-value) = 5.04) was found in Ssa05 in a region of 6,7 Kb, associated with the CR762469.1 gene; other high scores were found in Ssa10 and Ssa01, nearby to *mipol1, furinb, csnklg2a* and *rs17*. Other candidate genes undergoing selection for this population are shown in Supplementary Table S1.

**Figure 2.**
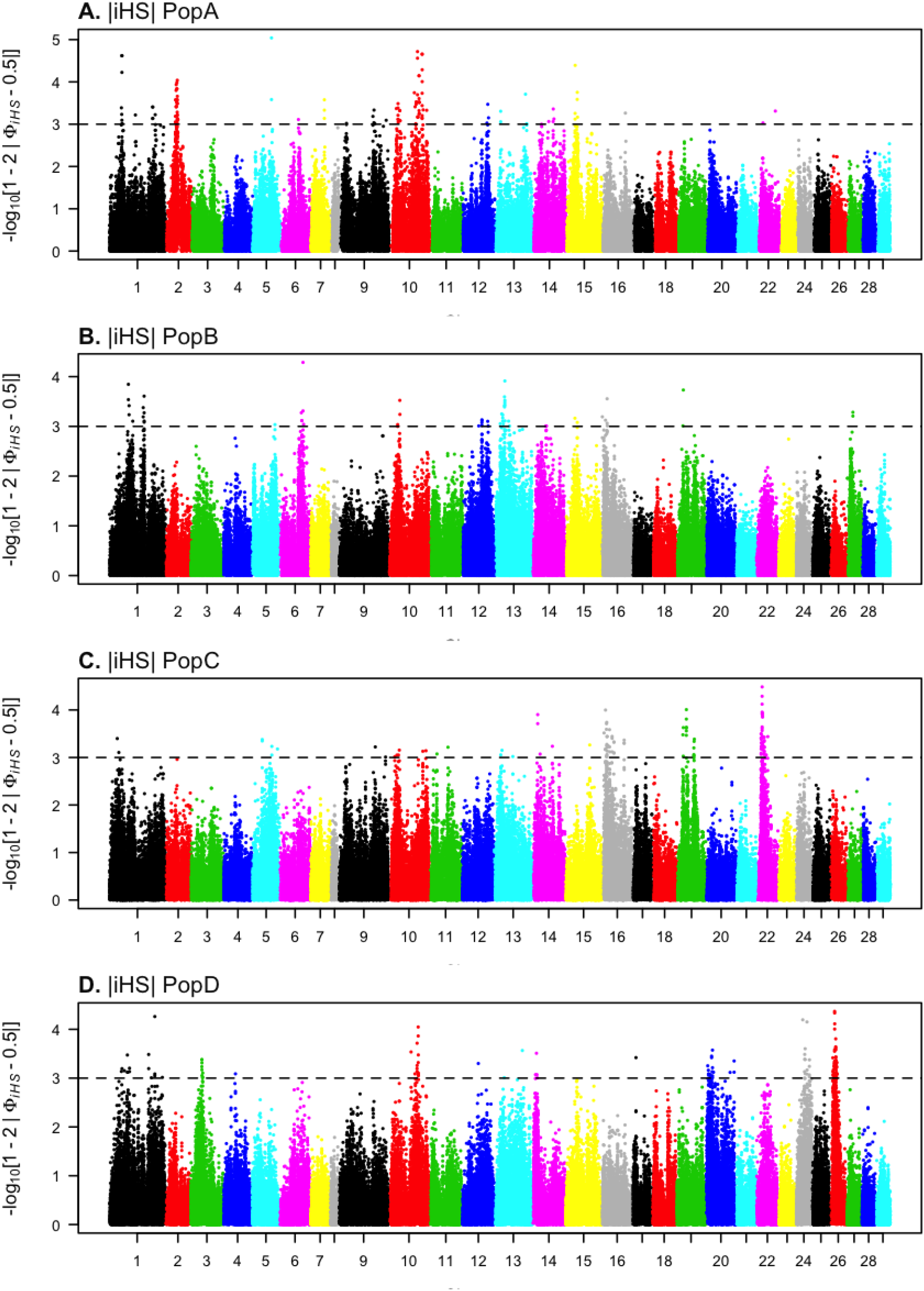
Genome-wide distribution of −log10(p-value) of standardized Integrated Haplotype Score |iHS| among Atlantic salmon populations.

**Table 2.** Number of SNPs identified by |iHS| among populations and chromosomes.

|  | Pop-A | Pop-B | Pop-C | Pop-D |
| --- | --- | --- | --- | --- |
| Ssa01 | 14 | 11 | 2 | 15 |
| Ssa02 | 31 |  |  |  |
| Ssa03 |  |  |  | 9 |
| Ssa04 |  |  |  | 1 |
| Ssa05 | 2 | 1 | 5 |  |
| Ssa06 | 1 | 6 |  |  |
| Ssa07 | 3 |  |  |  |
| Ssa08 |  |  |  |  |
| Ssa09 | 6 |  | 2 |  |
| Ssa10 | 43 | 3 | 6 | 14 |
| Ssa11 |  |  | 2 |  |
| Ssa12 | 3 | 7 |  | 1 |
| Ssa13 | 4 | 16 | 3 | 2 |
| Ssa14 | 4 | 3 | 4 | 4 |
| Ssa15 | 6 | 2 | 1 |  |
| Ssa16 | 1 | 5 | 32 |  |
| Ssa17 |  |  |  | 1 |
| Ssa18 |  |  |  |  |
| Ssa19 |  | 2 | 15 |  |
| Ssa20 |  |  |  | 18 |
| Ssa21 |  |  |  |  |
| Ssa22 | 2 |  | 49 |  |
| Ssa23 |  |  |  |  |
| Ssa24 |  |  |  | 15 |
| Ssa25 |  |  |  |  |
| Ssa26 |  |  |  | 54 |
| Ssa27 |  | 2 |  |  |
| Ssa28 |  |  |  |  |
| Ssa29 |  |  |  |  |
| Total | 120 | 58 | 121 | 134 |

In Pop-B fourteen regions passed the threshold, distributed among eight chromosomes (Ssa1, 6, 10, 12, 13, 14, 16, and 27). The highest score was in Ssa06, harboring the SASH1 gene. Ssa01 and Ssa13 encompassed 4 and 3 regions under selection, respectively, spanning from 11 Kb to 228 Kb. A total of 24 genes were located in these regions (Supplementary Table S1).

In Pop-C |iHS| detected 121 SNPs passing the threshold and we annotated sixteen genomic regions. Ssa22 showed the highest scores and larger regions under selection, harboring genes such kcnkf, *sc61a, mapk3, f264* and *cdh2*. Ssa16 and Ssa19 also exhibited high |iHS| scores spanning regions 3 Kb to 1788 Kb.

Finally, Pop-D presented the highest number of SNPs (134 SNPs) above the threshold compared with other populations, distributed across 11 chromosomes. We defined 25 genomic regions under selection, most of them located in Ssa26, where the highest |iHS| scores were also found. Genes such as uqcrfs1, neto1, itfg1 and phkb were found in these regions. Ssa24 also presented higher |iHS| values in one of its regions associated with tchp, ube3b and myo1ha among others. Details of genes and regions can be found in Supplementary Table S1.

### Candidate regions under selection – *XPEHH*

We also looked for selection signatures using the XPEHH test between the following populations pairs: A/B; A/C and A/D (Figure 3). We detected, 437 (A/B), 764 (A/C) and 262 (A/D) XPEHH scores outlier SNPs indicative of selection (Table 3). We considered potential genomic regions under selection as those containing two or more consecutive SNPs less than 500 Kb apart and that had XPEHH score > 3. After merging overlapping regions 27, 25 and 15 candidate regions were identified for A/B, A/C and A/D comparisons respectively. The total length of the candidate regions was 10.13 Mb for A/B, 12.11 Mb for A/C and 4.05 Mb for A/D. Comparison between A/C yielded negative results in chromosome Ssa14 and Ssa16, furthermore comparison A/D yielded negative results in Ssa14, Ssa24 and Ssa26, suggesting selection in the reference population (A). The gene annotation revealed in A/B the *plecb* gene on Ssa02 and *myo1cb, slc43a2a* and *ywhae1* genes on Ssa09, associated with the highest values of XPEHH. in A/C the chromosome Ssa10 presented a large number of SNPs and regions putatively under selection. Also this chromosome presented the highest scores; genes such as *fnbp1l, bcar3*, slc5a9 and *fryl*were associated with these values. The highest values for A/D were also located on Ssa10 with *lhx4*, *shr3rf1* and *ftr33* genes. The negative values of XPEHH harboring genes such *agla*, *kcmf1, cds1* and *tshz3b* suggest selection on population A.

**Figure 3.**
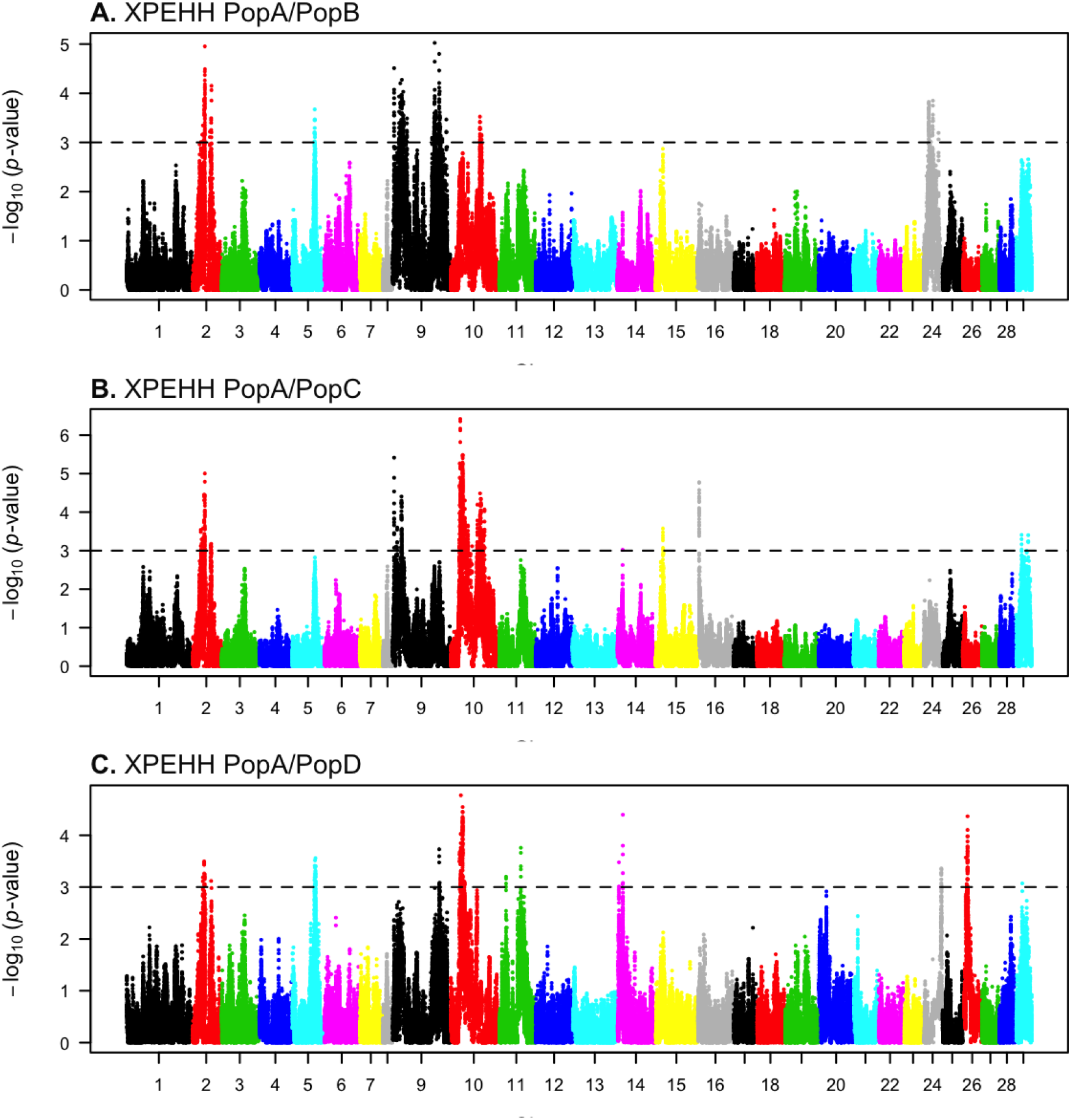
Genome-wide distribution of −log10(p-value) of standardized cross-population extended haplotype homozygosity (XP-EHH) scores in pairwise Atlantic salmon populations.

**Figure 4.**
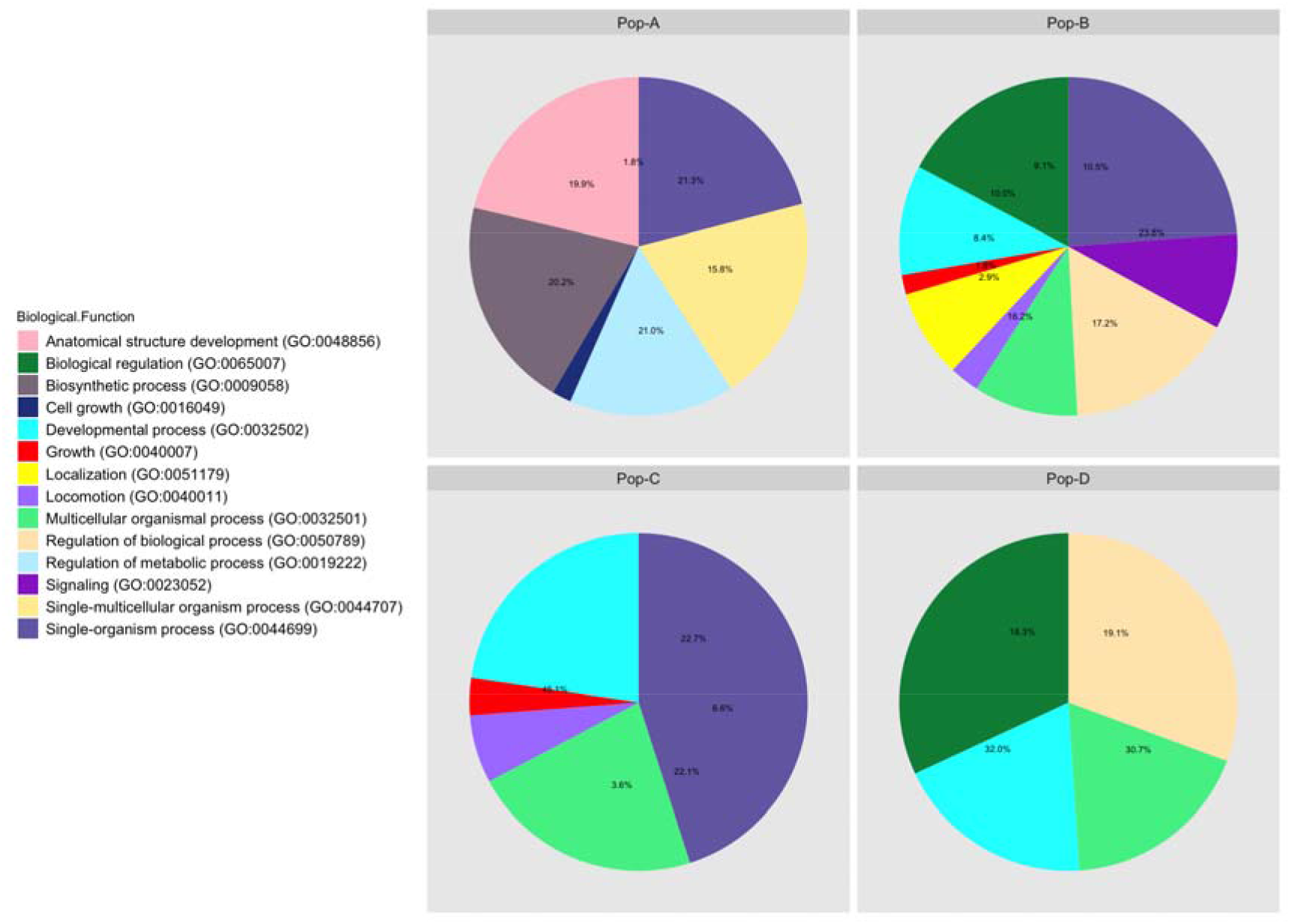
GO enrichment analysis of genes with evidence of selection in Atlantic salmon. GO functional classification was performed using the DAVID browser.

**Table 3.** SNPs and regions under selection identified by *XPEHH* among population pairs and chromosomes.

|  | Pop-A/Pop-B |  | Pop-A/Pop-C |  | Pop-A/Pop-D |  |
| --- | --- | --- | --- | --- | --- | --- |
|  | SNPs | Regions | SNPs | Regions | SNPs | Regions |
| Ssa01 |  |  |  |  |  |  |
| Ssa02 | 68 | 3 | 82 | 5 | 17 | 2 |
| Ssa03 |  |  |  |  |  |  |
| Ssa04 |  |  |  |  |  |  |
| Ssa05 | 11 | 1 |  |  | 26 | 3 |
| Ssa06 |  |  |  |  |  |  |
| Ssa07 |  |  |  |  |  |  |
| Ssa08 |  |  |  |  |  |  |
| Ssa09 | 289 | 17 | 71 | 3 | 6 | 1 |
| Ssa10 | 16 | 2 | 571 | 13 | 154 | 4 |
| Ssa11 |  |  |  |  | 10 | 2 |
| Ssa12 |  |  |  |  |  |  |
| Ssa13 |  |  |  |  |  |  |
| Ssa14 |  |  | 1 |  | 8 | 1 |
| Ssa15 |  |  | 7 | 1 |  |  |
| Ssa16 |  |  | 25 | 1 |  |  |
| Ssa17 |  |  |  |  |  |  |
| Ssa18 |  |  |  |  |  |  |
| Ssa19 |  |  |  |  |  |  |
| Ssa20 |  |  |  |  |  |  |
| Ssa21 |  |  |  |  |  |  |
| Ssa22 |  |  |  |  |  |  |
| Ssa23 |  |  |  |  |  |  |
| Ssa24 | 53 | 4 |  |  | 8 | 1 |
| Ssa25 |  |  |  |  |  |  |
| Ssa26 |  |  |  |  | 32 | 1 |
| Ssa27 |  |  |  |  |  |  |
| Ssa28 |  |  |  |  |  |  |
| Ssa29 |  |  | 7 | 2 | 1 |  |
| Total | 437 | 27 | 764 | 25 | 262 | 15 |

### Gene ontology for candidate genes under selection

To further explore the functions of the candidate genes nearby markers showing evidence of selection signatures, we annotated the candidate genes detected by both methods using DAVID browser (https://david-d.ncifcrf.gov). These candidate genes were enriched in 14 gene ontology terms. None of these categories were common across all four populations, but Developmental process and Multicellular organismal process were common on Pop B-C and D. Regulation of biological process was shared for Pop A –B and D; Single-organism process was common for Pop A –B and C; Biological regulation was found in Pop-B and Pop-D; and Growth and Locomotion were common for Pop-B and Pop-C. Anatomical structure development, Biosynthetic process, Cell growth and Single-multicellular organism process were present only in Pop-A; while Localization and Signaling were found only in Pop-B (4).

## 5. DISCUSSION

In this study two complementary tests were used to detect genome wide selection signatures within and between four Atlantic salmon populations with Norwegian origin. We used |iHS| test to evaluate selection signatures within populations and XPEHH to evaluate across populations. We used the oldest population as the reference population when using XPEHH to evaluate the effect of domestication and artificial selection in three different locations in Chile.

### Structure and diversity

To examine genetic population structure and relationships among the major groups of salmon, we conducted an ADMIXTURE analyses based on high-quality SNP data. This analysis revealed twelve clusters, which was expected considering the admixed origin of these populations (Verspoor *et al*. 2007). The four populations used in this study come from the Mowi strain, which was created, using samples from several rivers along the west coast of Norway (Norris *et al*. 1999). The population with the lowest level of admixture was Pop-A, which was also the population with the lowest genetic diversity, a condition that could reflect a higher intensity of artificial selection in this population. Intense artificial selection causes loss of genetic variation as a consequence of the mating of related individuals (Gjedrem 2005). Pop-B and Pop-C showed very similar patterns of heterozygosity and admixture level, which was expected due to the similar breeding practices and environmental conditions to which they have been subjected. Pop-D, however, showed the highest level of heterozygosity and a more complex pattern of admixture, likely produced by a lower pressure of artificial selection on this population. Recent genetic introgression cannot be discarded for Pop-D given the potential of crosses with a different strain for management issues. The results presented here also reinforce the notion that a few generations (at least four in this particular case) are sufficient to generate large changes in terms of genetic structure in farmed Atlantic salmon populations, with the same genetic origin, which have been subjected to different management and environmental conditions. Estimates of inbreeding coefficient (F_IS_) showed the lowest value in Pop-D, which is consistent with the heterozygosity level in this population. Pop-A presented the second lowest value, despite the fact that this population has been subjected to the most intense selection pressure probably due to a better inbreeding management with the use of DNA fingerprinting technology to know relatedness among individuals in order to avoid inbreeding.

### Selection signatures

As expected the highest |iHS| scores were found in Pop-A because this population has been subjected to more intense artificial selection pressures for a longer time. The number of SNPs under selection detected by this method was similar in Pop-A, Pop-C and Pop-D, but lower in Pop-B. We suggest that this difference is due to the fact that the |iHS| test has little power to detect signals near fixation (Sabeti *et al*. 2007; Simianer *et al*. 2010). XPEHH, which is more powerful at detecting selection signatures at or near fixation (Sabeti 2007), detected a similar number of regions putatively under selection in Pop-B and Pop-C, but lower in Pop-D. Conversely, |iHS| detected more SNP in this population, suggesting loci under selection in Pop-D have experienced weaker pressure of artificial selection and a greater impact of natural selection, which has prevented allele fixation. Overlaps among regions detected by |iHS| method, were found only when using pairs of populations, that is, a common region was found between Pop-A/B, Pop-A/C, Pop-A/D, Pop-B/C, Pop-B/D and Pop-C/D. No overlap was found among four populations or when using any combination of three. XPEHH detected a higher number of shared regions among populations, specifically in Ssa02, which was common to all 3 tested populations. In addition, shared regions were found in population pairs B/C and Pop-D/C. A greater number of shared regions detected by XPEHH could be explained by a greater power to detect regions that have experienced older selection events (Sabeti *et al*. 2007; Klimentidis *et al*. 2011) than those detectable by |iHS|. Therefore, these regions may be explaining selection signatures that originated before these populations were brought to Chile.

### Domestication traits in salmon

Selection signatures found in this study may be involved in some desirable economic traits in salmon production as well as traits that are typically under the effect of domestication. All populations used in this study have been subjected to artificial selection to improve growth rate. According to the functional annotations of the candidate genes, several biological processes were found to be involved with growth and development, such as the Development process and Regulation of Biological/metabolic processes. Additionally, some of the genes identified have been associated with growth traits in other species; such *sh3rf1* in chicken and cattle (Hanotte *et al*. 2003; Rubin *et al*. 2010) or *igf1r*, which was previously found to be a size locus of large effect in dogs (Sutter *et al*. 2007; Hoopes *et al*. 2012). We suggest that these genes may be under selection for improving growth related traits in salmon. On the other hand, we also identified genes such *scaper, clstn3* and *pex5* related to mental disorders in humans (Glatt *et al*. 2005; Pettem *et al*. 2013). Other genes related to behavioral traits have be found in other Atlantic salmon strains, as well (Lopez et al, 2018), suggesting that artificial selection acts on behavioral traits in salmon as in other domestic animals (Clutton-Brock 1999).

## 6. CONCLUSIONS

In the present study, several candidate genomic regions with selection signatures were identified using two haplotype based methods, |iHS| and XPEHH in four populations of Atlantic salmon. These genomic regions harbored important genes that enriched G terms including growth, developmental processes, and have been associated with growth and behavior in other species. These finding improve our understanding of genomic variants undergoing selection in domestic populations of Atlantic salmon.

## Ethics approval and consent to participate

The sampling protocol was previously approved by The Comité de Bioética Animal, Facultad de Ciencias Veterinarias y Pecuarias, Universidad de Chile (Certificate N° 29– 2014).

## Consent for publication

Not applicable

## Availability of data and material

Genotype data for each population is available from the online digital repository *figshare* https://figshare.com/s/83efa70722ed5ada023a

## Author’s contributions

MEL and JMY conceived the research idea. MEL drafted the manuscript and carried out the analyses. TL supervised the data analyses and contributed to discussion and writing. TL, AN, JPL, RN, and JMY reviewed the manuscript. All authors read and approved the final manuscript.

## Acknowledgements

MEL acknowledges the National Commission of Scientific and Technologic Research (CONICYT) for the funding through the National PhD funding program. JMY is supported by Núcleo Milenio INVASAL funded by Chile’s government program, Iniciativa Científica Milenio from Ministerio de Economía, Fomento y Turismo

